# Peptidylarginine Deiminase inhibition abolishes the production of large extracellular vesicles from *Giardia intestinalis*, affecting host-pathogen interactions by hindering adhesion to host cells

**DOI:** 10.1101/586438

**Authors:** Bruno Gavinho, Izadora Volpato Rossi, Ingrid Evans-Osses, Sigrun Lange, Marcel Ivan Ramirez

## Abstract

*Giardia intestinalis* is an anaerobic protozoan that is an important etiologic agent of inflammation-driven diarrhea worldwide. Although self-limiting, a deep understanding of the factors involved in the pathogenicity that produces the disruption of the intestinal barrier remains unknown. There is evidence that under diverse conditions, the parasite is capable of shedding extracellular vesicles (EVs) which could modulate the physiopathology of giardiasis. Here we describe new insights of *G. intestinalis* EV production, revealing its capacity to shed two different enriched EV populations (large and small extracellular vesicles) and identified a relevant adhesion function associated only with the larger population. Our work also aimed at assessing the influences of two recently identified inhibitors of EV release in mammalian cells, namely peptidylarginine deiminase (PAD) inhibitor and cannabidiol (CBD), on EV release from *Giardia* and their putative effects on host-pathogen interactions. PAD-inhibitor Cl-amidine and CBD were both able to effectively reduce EV shedding, the PAD-inhibitor specifically affecting the release of large extracellular vesicles and interfering with *in vitro* host-pathogen interactions. The strong efficacy of the PAD-inhibitor on *Giardia* EV release indicates a phylogenetically conserved pathway of PAD-mediated EV release, most likely affecting the *Giardia* arginine deiminase (GiADI) homolog of mammalian PADs. While there is still much to learn about *G. intestinalis* interaction with its host, our results suggest that large and small EVs may be differently involved in protozoa communication, and that EV-inhibitor treatment may be a novel strategy for recurrent giardiasis treatment.

## Introduction

*Giardia intestinalis* is a lumen dwelling pathogen in the vertebrate gut, responsible for a worldwide waterborne diarrhea known as giardiasis. The flagellated protozoa was formerly incorporated in the WHO neglected diseases, estimated a burden not only for poor, but also industrialized countries (Savioli et al., 2006). Approximately 300 million infections are identified annually, mainly in children (reviewed by Ankarklev et al., 2010; Cernikova et al., 2018). Its life cycle consists of two evolutionary stages: i) the trophozoite, which adheres to the intestinal mucosal barrier and multiplies by binary fission, and ii) the infectious stage, the cyst, that is released through feces and is acquired by ingestion of food or water,. Although not invasive, this extracellular protozoa has a distinct cellular structure due to evolutionary reduction (Cernikova et al., 2018). *G. intestinalis* interaction with the host, and its immune evasion, are mediated through many survival factors, including inhibition of neutrophil migration through cathepsin B (Cotton et al., 2015), arginine and cytokine suppression by arginine deiminases (Touz et al., 2018), induction of epithelial translocation through cysteine proteases (Liu et al., 2018), anti-oxidant production, microvillus shortening (reviewed by Bartelt & Sartor, 2015), dysbiosis (Barash et al., 2017; Bartelt et al., 2017) and antigenic variation (Prucca et al., 2008; Serradel et al., 2016). A complete comprehension of the pathogenesis is still elusive, and many factors are involved in the persistence of trophozoites on the host, including drug resistance (Serradell et al., 2016). Finally, chronic infection is a significant concern in giardiasis, due to sequelae related to nutritional and cognitive deficiency in children and immunocompromised individuals (Halliez & Buret, 2013; Bartelt & Sartor, 2015). Adaptions of the parasite for survival in the host involve sophisticated forms of host-pathogen communication. Many reports have described the release of extracellular vesicles (EVs) from pathogens to be relevant to disease status (Cwiklinski et al., 2015; Coakley et al., 2017). EVs are found in most biological fluids and are 30-1000 nm lipid-bilayer vesicles, which are shed from cells and transport a range of biomolecules, participating in cell communication in physiological and pathophysiological processes (Coakley et al., 2015; Maas et al., 2017; van Niel et al., 2018; Ramirez et al., 2018; Ryu et al., 2018). Our group has previously described EV release of *G. intestinalis* and established that protozoa EVs are involved in pathogen interactions via immunomodulation and trophozoite persistence (Evans-Osses et al., 2017). In recent years, there has been a growing interest to improve current knowledge of the nature, constitution and biogenesis of all secreted EVs. Lately, the field of EVs is debating the need to accurately separate EV subtypes to investigate functional relevance (Tckach et al., 2018; Slomka et al., 2018). Enrichment of subpopulations of EVs from samples is acquired based on differential centrifugation steps: large EVs (LEVs) are obtained at speeds lower than 20,000xg and small EVs (SEVs) are pelleted at 100.000xg in a further ultracentrifugation step. Formerly, microvesicles and exosomes, particularly present in LEV and SEV respectively, are considered the main EV subpopulations. Exosomes are the smallest population, continually produced in the late endosome and liberated through the fusion of multivesicular bodies within the plasma membrane. Microvesicles are particles of a bigger size produced though a budding from the plasma membrane under stress mediated by scramblase, calpain and Ca^2+^ liberation (Morrison et al., 2016). Different roles for these EV subpopulations remain a focus of ongoing investigations; are all EVs phenotypically relevant and/or similar? Another aspect of great importance is EV cargo and its modulation during biological processes or via drug-treatment. There is growing evidence that nucleic acids are also found in protozoa EVs, including dsDNA, tRNA, rRNA and small RNAs that could modulate gene expression from recipient cells (Kim et al., 2017; Tsatsaronis et al., 2018). The identification of the involvement of nucleic acids generated by pathogen or host cells, and transferred through EVs, have initiated a new platform for investigation in parasitology (Coakley et al., 2015).

While it is known that EVs are released by multiple mechanisms, some advances in understanding of their biogenesis has been elucidated via studies on the peptidylarginine deiminase (PAD)-mediated pathway of EV release (Kholia et al., 2015; Kosgodage et al., 2017; Lange et al., 2017; Kosgodage et al., 2018a). PADs are phylogenetically conserved enzymes from bacteria to mammals (Vossenaar et al., 2003; Magnadottir et al., 2018), including in *Giardia* (arginine deiminase GiADI; Trejo-Soto et al., 2016). PADs catalyze post-translational deimination by irreversibly changing arginine into citrulline in a calcium-catalyzed manner in target proteins, affecting their folding and function (Vossenaar et al., 2003; György et al., 2006). PADs are involved in physiological and pathophysiological processes and their upregulation and associated increase in deiminated proteins is associated with various pathologies including autoimmune and neurodegenerative diseases as well as cancer (Vossenaar et al., 2003; Wang and Wang 2013; Witalison et al., 2015; Lange et al., 2017). While exact roles for PADs in EV biogenesis and release remain to be fully elucidated, effects on cytoskeletal, nuclear and mitochondrial proteins have been reported (Kholia et al., 2015; Kosgodage et al., 2018). As pharmacological PAD-inhibitors have previously been shown to be potent inhibitors of EV release in various cancer cells, as well as to modulate EV cargo (Kholia et al., 2015; Kosgodage et al., 2017; Kosgodage et al., 2018a), we sought to investigate a phylogenetically conserved influence of such PAD-inhibitors on the EV production of our protozoa model; putatively targeting GiADI and therefore also reveal a role for GiADI in EV release.

In addition, cannabidiol (CBD), a phytocannabinoid derived from *Cannabis sativa* (Mechoulam et al., 2002) was recently identified as a potent EV-inhibitor in cancer cells (Kosgodage et al., 2018b, Kosgodage et al., 2019). As cannabinoids have previously been associated with anti-parasitic functions (Nok et al., 1994, Croxford et al., 2005; Roulette et al., 2016) and immunoregulatory roles during infectious disease (reviewed in Hernández-Cervantes et al., 2017) we sought to identify whether EV release from *Giardia* may be affected by CBD, thus elucidating a novel aspect of CBD function on *Giardia*-host interaction.

Here, we report that *G. intestinalis* produces two populations of EVs that differ in size. The larger EV population had a significant effect on protozoa-host adhesion *in vitro* and was significantly reduced by both PAD-inhibitor and CBD. In addition, treatment with PAD-inhibitor selectively prevented protozoa EV production.

## Methods

### *G. intestinalis* isolates and cell culture

*G. intestinalis* isolate WB (ATCC 50803) were grown in TYI-S-33 medium (Keister, 1983) supplemented with 10% adult bovine serum with 1% Penicillin/Streptomycin 1000 U (Gibco ™) and 0.05% bovine bile (ThermoFisher ™) at 37°C under microaerophilic conditions. The cultures were maintained until confluent and thereafter sub-cultured, each for 72 hours. Human colorectal adenocarcinoma cells, caco-2 (ATCC CRL-2102) were cultured in RPMI supplemented with 10% fetal calf serum and 1% Penicillin/Streptomycin 1000 U (Gibco™). Cells were incubated at 37°C, 5% CO_2_ until a confluent cell monolayer was reached.

### EV isolation

Parasites from confluent cultures were decanted by chilling for 15 min in ice-cold, centrifuged twice (600xg/5min) and the pellets suspended with fresh TYI-S-33 without adult bovine serum (FBS). Parasites were then counted using a hemocytometer, and diluted to 1×10^6^ per sample according to Evans-Osses et al.(2017). 1mM of CaCl_2_ were added to the parasite culture for EV induction and incubated for 1 hour at 37°C. Then, EV pellets were collected via step-wise centrifugation: first, at 600xg/5 min; 4000xg/30 min to eliminate cellular debris, thereafter the supernatant was centrifuged at 15,000xg for 1h and the resulting pellet was washed and resuspended in phosphate buffered saline (PBS). The remaining supernatant was then ultracentrifuged for 100,000xg for 4h, and the resulting EV-containing pellets were washed once in PBS. Both samples were kept at 4°C until further use. To determine cell viability post-EV isolation, parasites were adjusted to 1×10^6^ per sample and submitted to the vesiculation protocol for 1, 3 or 6 hours. Then, trophozoite pellets were subcultured. For mammalian EV purification, caco-2 were cultured until confluence, and then medium was removed. Cultures were washed twice with fresh RPMI-1640 and kept for 1 h with medium without FBS. The supernatant was processed in the same manner as for protozoa.

### EV Quantification and Characterization

EVs were in the first instance quantified based on their protein concentrations by the Micro BCA assay (ThermoFisher ™). For Nanoparticle tracking analysis (NTA, Nanosight, Malvern, U.K.), each sample was diluted 1:100 in PBS and subjected to a NS300 Nanosight, with readings performed in triplicate during 60 s videos at 10 frames per second at room temperature, with the following parameters: camera shutter – 1492, camera gain – 512, detection threshold – 10. The resulting replicate histograms were averaged for presentation in box-plots.

### Treatment of trophozoites with EV-inhibitors

Inoculum of 10^6^ trophozoites per group (triplicates) were stimulated with1 mM CaCl_2_ for the production of EVs in microtubes with or without EV-inhibitors. Groups were as follows: medium only (control), 100 or 50 μM PAD-inhibitor Cl-amidine (a kind gift from Prof Paul Thompson, UMASS), or with 10 or 5 μM CBD (90899_SIAL, Sigma-Aldrich™). After 60 minutes of incubation (37°C), samples were processed according to Evans-Osses et al (2017).

### Host-pathogen interaction assay after exposure with EV-inhibitors

caco-2 were seeded in 24-well plates and grown to 100% confluence. Inoculations of 5×10^5^ trophozoites per group were submitted to the EV production and then transferred to the cell monolayer for 3 hours (37°C) in a final volume of 1 mL / well (MOI 10:1). The following groups were investigated: medium only (control), 10 μM CBD, 10 μM CBD + 14 μg EVs, 100 μM Cl-amidine, 100 μM Cl-amidine + 14 μg EVs. After incubation, the trophozoites quantification was performed after removal of the supernatant, centrifuging non-adherent parasites and counting them using a hematocytometer. The percentage of trophozoites adhering to caco-2 cells was subsequently calculated according to Cotton et al. (2015). In experiments assessing the effects of mammalian EVs on trophozoite adhesion, caco-2 cells were grown until confluence, then the medium was removed and wells were washed and maintained with medium without FBS. Monolayers were thereafter treated with 100 µM Cl-amidine, for 1 h, followed by incubation with 5×10^5^ trophozoites per well. Experimental group were as follows: 7 or 14 µg caco-2 SEVs, 7 or 14 µg caco-2 LEVs. Statistical analysis was performed by one-way ANOVA (*p*<0.0001).

### EV staining

For uptake assays, EVs were stained and tested with carboxyfluorescein succinimidyl ester (CFSE, ThermoFisher™) or with the lipophilic dye PKH-26 (SigmaAldrich™). For CFSE labelling, 1 µL of the fluorescent dye was diluted with both EV populations in 1 mL PBS. For PKH-26, 2 µL of the fluorescent dye were diluted in 1 mL of diluent C and both EV populations were diluted 1/40 in diluent C. Both dilutions were mixed together at a volume ratio of 1:1. For both fluorescent dyes, labelling was continued for 15 min at room temperature in the dark. The reaction was stopped by adding 1 mL FBS, and samples were then washed in PBS, and ultracentrifuged at at 15.000xg for 1 h to obtain LEVs and at 100,000xg for 4 h for collection of SEV.

### Protozoan EVs uptake by caco-2 cells

Caco-2 cells were incubated on sterile coverslips at 37°C in 5% CO_2_ with 3.5 or 14 μg of PKH26-labeled protozoal EVs (ThermoFisher™) for 1h. Caco-2 monolayers were also labelled for nuclei (DAPI, blue - ThermoFisher™). After incubation, the cells were extensively washed in cold PBS, and fixed with 4 % paraformaldehyde. Coverslips were washed with PBS and mounted with 10 μl of a 50 % glycerol solution. Internalized EVs were detected by confocal microscopy (Nikon A1R HD Multifoton Confocal). Images were processed by Image J software (v. 1.48 – open source, Schneider et al., 2012). Fluorescence intensity of two images per sample were obtained in a duplicate experiment, and corrected cellular fluorescence was estimated as in McCloy et al (2014).

### Cytotoxicity Assay of EV-inhibitors toward caco-2 cells

Caco-2 were seeded into a 96-well plate and grown at 37°C in 5% CO_2_ until confluence in RPMI-1640 medium containing 10% fetal bovine serum and 1% penicillin/streptomycin (10 000 UI). Cells were treated with albendazole (ABZ) at 10 μM, 100 μM Cl-amidine and 10 μM CBD, and the volume was adjusted to 100 μL of RPMI per well. After 48 hours, wells were washed with 100 μL of PBS (50%). Cells were fixed with methanol (50 μL) for 10 min, after which 50 μL of crystal violet 0.2% in ethanol/water (2% V/V) were added to each tube. After 2 min, the wells were exhaustively washed with 200 μL of PBS. Elution was made with a sodium citrate solution (0.05 μmol, 10 min), and absorbance was determined at 540 nm on a plate spectrophotometer.

## Results

### Giardia EV Biogenesis: identification of two distinct populations

The first objective was to obtain different EV populations from *G. intestinalis.* The previous protocol (Evans-Osses et al., 2017) was slightly modified to separate putative large extracellular (LEV) and small extracellular vesicles (SEV) from the total extracellular vesicles described before (Fig.1A). Two different EV populations were recovered from this method: LEVs at 15,000xg and SEVs at 100,000xg. The protocol was performed for protozoa and mammalian host cells. Protein estimation (Fig.1B) and nanoparticle tracking analysis (Fig.1C) showed a higher yield of protozoa EVs than caco-2 (∼3 fold higher). In addition, *Giardia* is capable of shedding more LEVs (∼ 3 fold higher than SEVs). LEV and SEV fractions showed a respective mean vesicle diameter of 187.6 and 67.7 nm respectively (Fig.1E-F). We also investigated cell viability after stimulating trophozoites to produce EVs for 1, 3 and 6 hours respectively, following the growth curve in a complete medium by 72 h at 37°C, 5%CO_2_. Parasites stimulated for 1 h-EV production maintained a normal growth (Fig. 1D). Parasites stimulated for EV production for 3 and 6 h only reached confluence after more than 72 h, therefore not maintaining normal growth rate.

**Figure 1.**
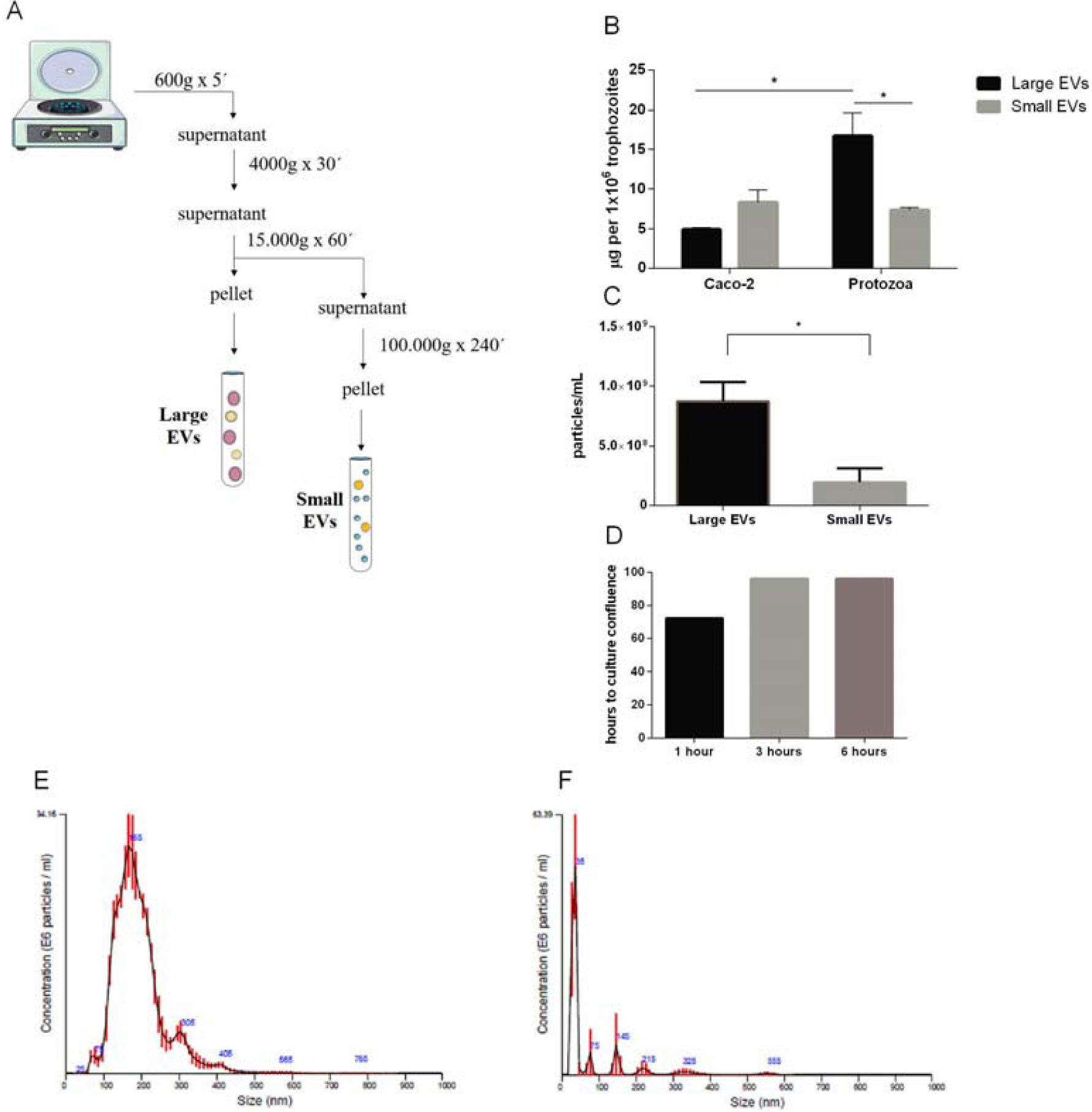
Isolation and characterization of two distinct EV populations produced from *Giardia intestinalis.* **A.** Protocol for the isolation of LEVs and SEVs based on differential centrifugation. **B.** Micro BCA protein estimation for the distinct EVs. **C.** Quantification of vesicle numbers from nanoparticle tracking analysis. **D**. Time-course for culture confluence of trophozoite induced to produce EVs for 1, 3 and 6 hours respectively. **E-F.** Particle size estimated by nanoparticle tracking analysis for LEVs **(E)** and SEVs **(F).**

### PAD-inhibitor and CBD treatment affects Giardia EV biogenesis

*G. intestinalis* trophozoites were treated with PAD-inhibitor Cl-amidine or CBD respectively to investigate their ability to inhibit EV production. Both compounds were able to significantly reduce production of EVs (Fig. 2A). In addition, we assessed whether it would be possible to block host-pathogen interactions following treatment with the EV-inhibitors. Indeed, both compounds were capable of decreasing trophozoites adhesion to the caco-2 monolayer through inhibition of EV release. In comparison, groups that were also treated with purified EVs had a higher adherence estimation (Fig. 2B).

**Figure 2.**
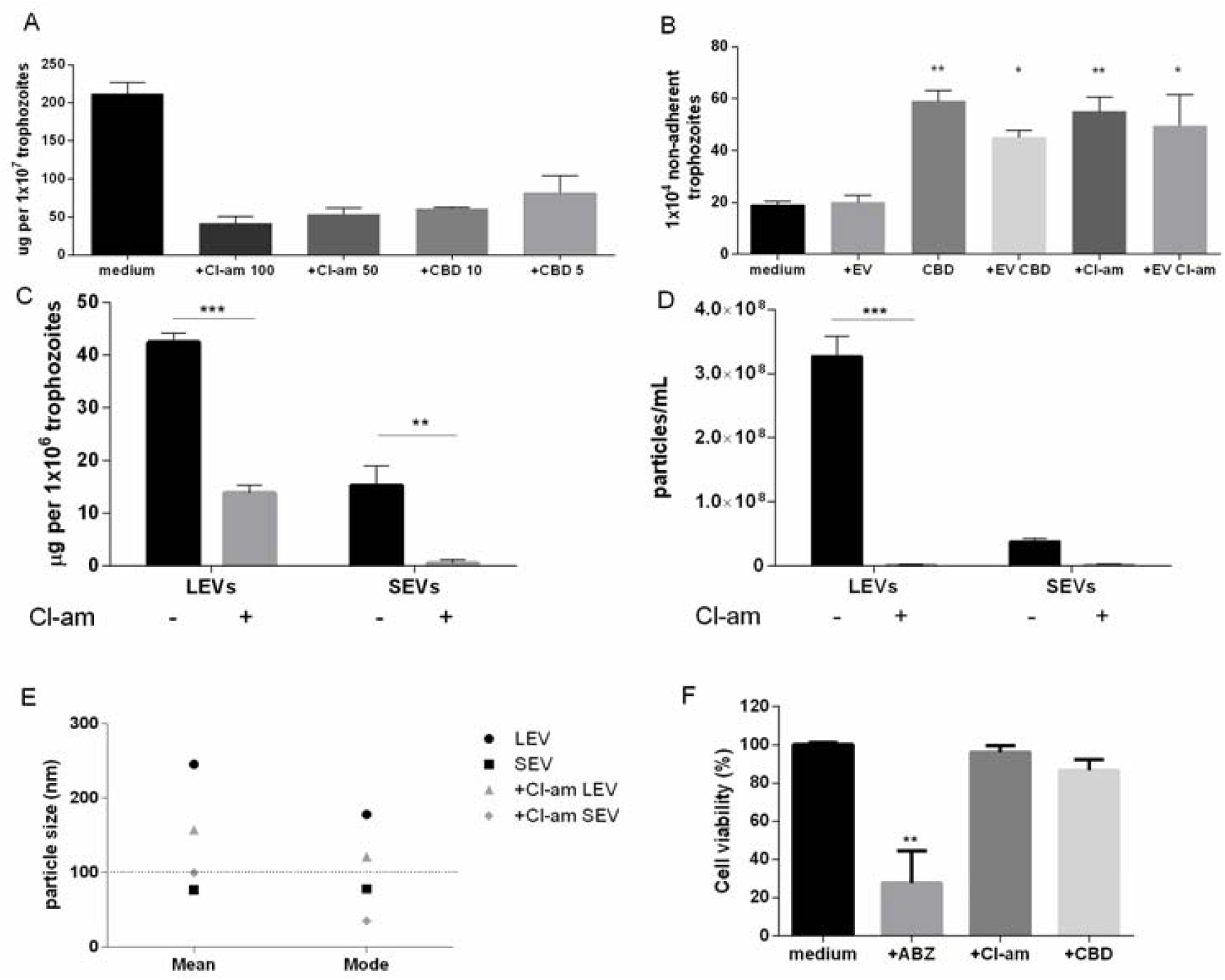
EV-inhibitors Cl-amidine and CBD decrease the production of LEVs secreted from *G. intestinalis.* **A.** EV protein quantification post-treatment with Cl-amidine (Cl-am; 100 and 50 µM) or CBD (10 and 5 µM). **B.** Adhesion assay post-treatment with PAD-inhibitor (100 µM) or CBD (10 µM). **C.** Protein estimation for the distinct EV populations following Cl-am treatment (100 µM) **D.** EV concentration estimation by nanoparticle tracking analysis (NTA) post-treatment with 100 µM Cl-am. **E.** EV size estimation of the distinct EV populations by NTA. **F.** Cytotoxic effects of the EV-inhibitors on caco-2 monolayers. Cells were incubated for 48 h with ABZ (albenzadole), Cl-am and CBD at 10, 100, and 10 μM respectively, or with culture medium. Cell viability was determined by the crystal violet method.

On this basis, and as Cl-amidine was found to be the strongest EV-inhibitor, we further assessed if Cl-amidine affected the release of SEVs and LEVs equally. Protein estimation (Fig.2C) showed that treatment with 100 µM Cl-amidine reduced the production of both EV types. However, concentration of LEVs by NTA estimated a significant difference between treated compared to non-treated groups (∼100 fold higher), while there was no difference observed for SEVs between treated versus control non-treated parasites (Fig.2D). The mean diameter (nm) of vesicles released in the presence of Cl-amidine was as follows: LEV (245.5), + Cl-am LEV (157.3), SEV (77.2), +Cl-am SEV (99.8) (Fig. 2E). Toxicity of caco-2 cells upon treatment with the EV-inhibitors was insignificant in 48 hours-interval compared to the positive control albenzadole, which is one of the reference drugs for Giardiasis treatment (Fig.2F).

### Host-pathogen interactions: both EV types are internalized by mammalian cells

SEVs and LEVs were analyzed for their ability to interact with host cells. Among the fluorochromes tested for EV labelling only PKH-26 showed a homogeneous staining. Both PKH26-labelled LEVs and SEVs were incubated with caco-2 monolayers for 1 hour at two concentrations (7 or 14 µg). Confocal microscopy revealed punctuated patterns of fluorescence distributed intracellularly (Fig.3A). Both populations appeared to be taken up by the host cells in a dose-dependent manner. LEV intensity internalization was overall observed to be higher, which may relate to larger vesicle size probably due to the bigger size of the particle (Fig. 3B)

**Figure 3.**
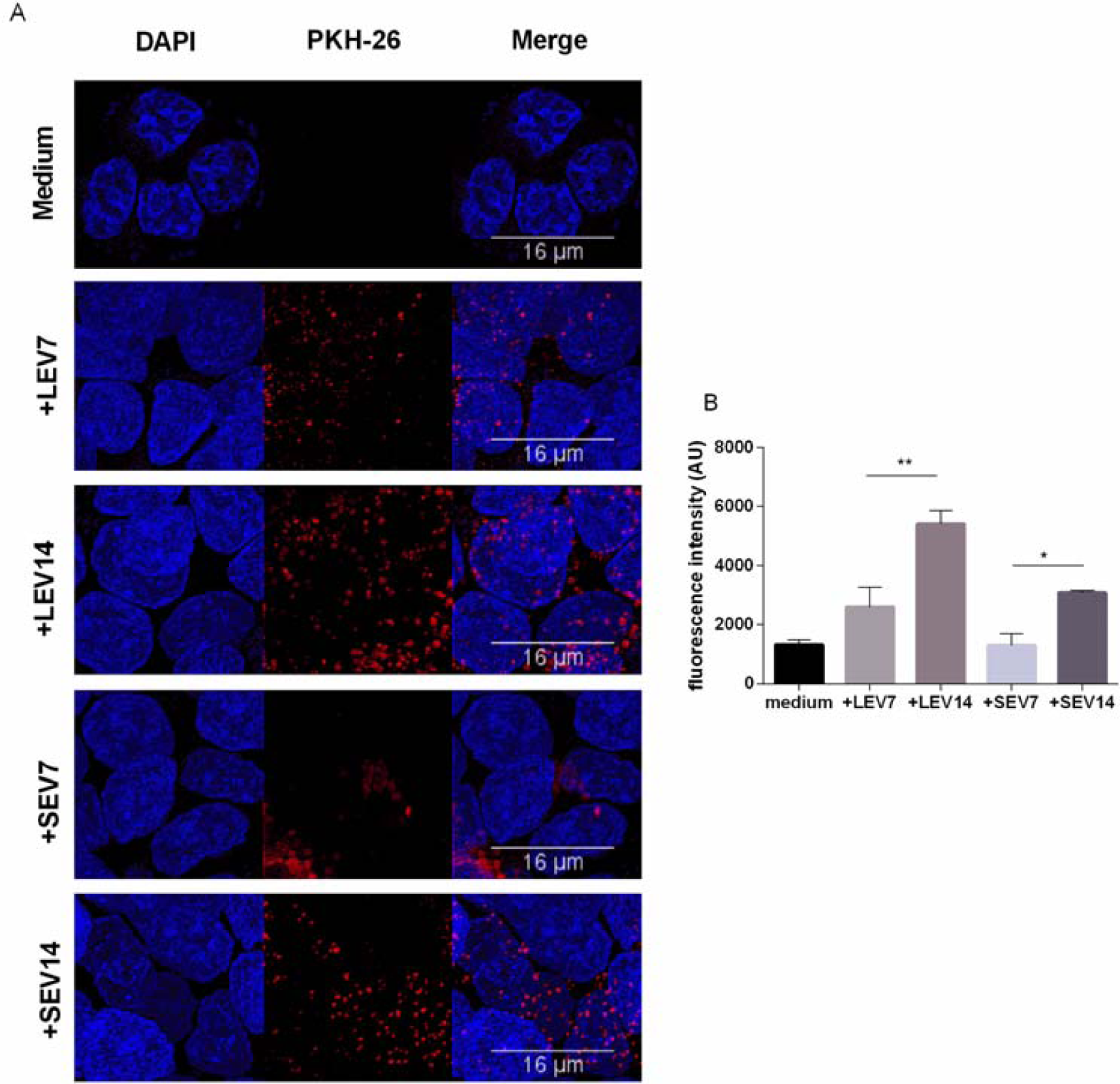
Both *G. intestinalis* EVs populations are efficiently taken up by caco-2. **A.** Caco-2 incubated with PKH26-labelled LEVs or SEVs. **B.** Internalized EVs were quantified by means of fluorescence intensity. Background signal was subtracted for every single image before obtaining the fluorescence intensity (Arbitrary Units); scale bars are indicated at 16 µm.

### LEVs derived from protozoa, but not host EVs restore the lack of adhesion to host cell of *G. intestinalis* trophozoites treated with EV-inhibitors

Due to the identification of two EV populations, LEVs and SEVs, in *G. intestinalis*, we sought to verify if each EV population have the same phenotype effect on host cell adhesion. For this assay, two concentrations (7 or 14 µg) from both EV subpopulations were used. Interestingly, LEVs derived from the protozoa were capable of restoring the adherence phenotype following treatment with Cl-amidine, in a dose-dependent manner (Fig.4A). In contrast, no effect was observed in the SEVs treated groups. These results suggest that physical properties related to adherence can be found in the larger *Giardia* EVs, and therefore EVs produced by the protozoa can selectively influence its phenotype.

Since a protozoa interaction with its host is complex and mediated through an active process, where it has been demonstrated that *Giardia* EVs participate in the adherence process, we also investigated if the host’s EVs could contribute to this phenomenon. Caco-2 monolayers were washed and treated with Cl-amidine, and trophozoites were added to the wells, followed by treatment with caco-2 EVs (Figure 4B). Opposed to what observed for the trophozoite EVs, mammalian LEVs or SEVs had no phenotypical effect on trophozoite adhesion.

**Figure 4.**
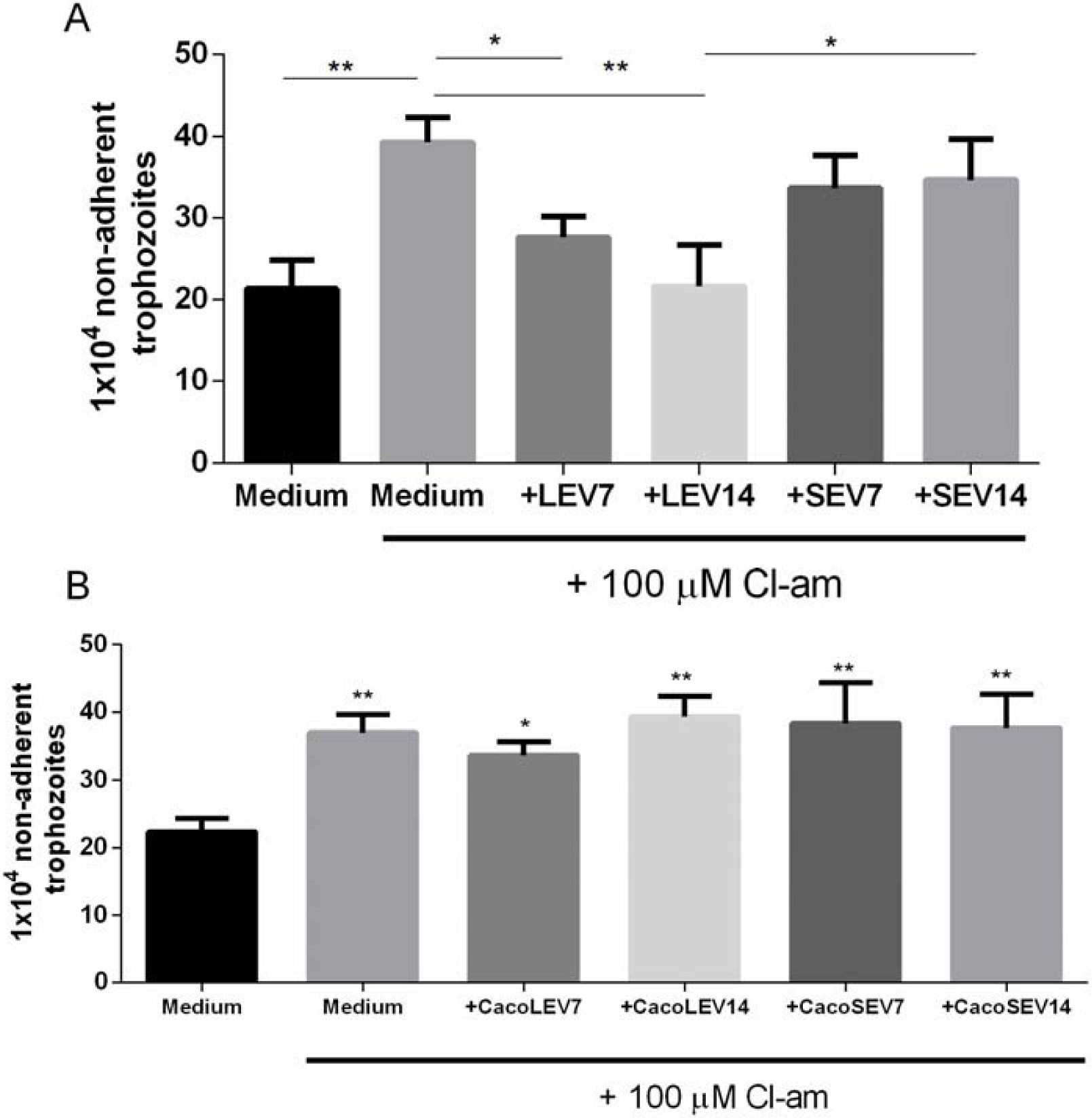
Protozoa derived EVs are selectively involved with protozoa adhesion to host cells. **A.** Adhesion assay following treatment with distinct protozoa EV populations and 100 µM Cl-amidine. **B.** Host-pathogen assay after treatment of caco-2 monolayer with Cl-amidine and incubation with mammalian cell derived EVs.

## Discussion

In this study, we describe two distinct EV populations from *Giardia intestinalis* where large EVs (LEVs), but not small EVs (SEVs), are associated with effective parasite cell adhesion to the host (Fig.5). Differing roles for these two EV populations, including in host-pathogen interactions, were demonstrated as treatment *of G. intestinalis* trophozoites with EV-inhibitors selectively decreased biogenesis of LEVs.

**Figure 5.**
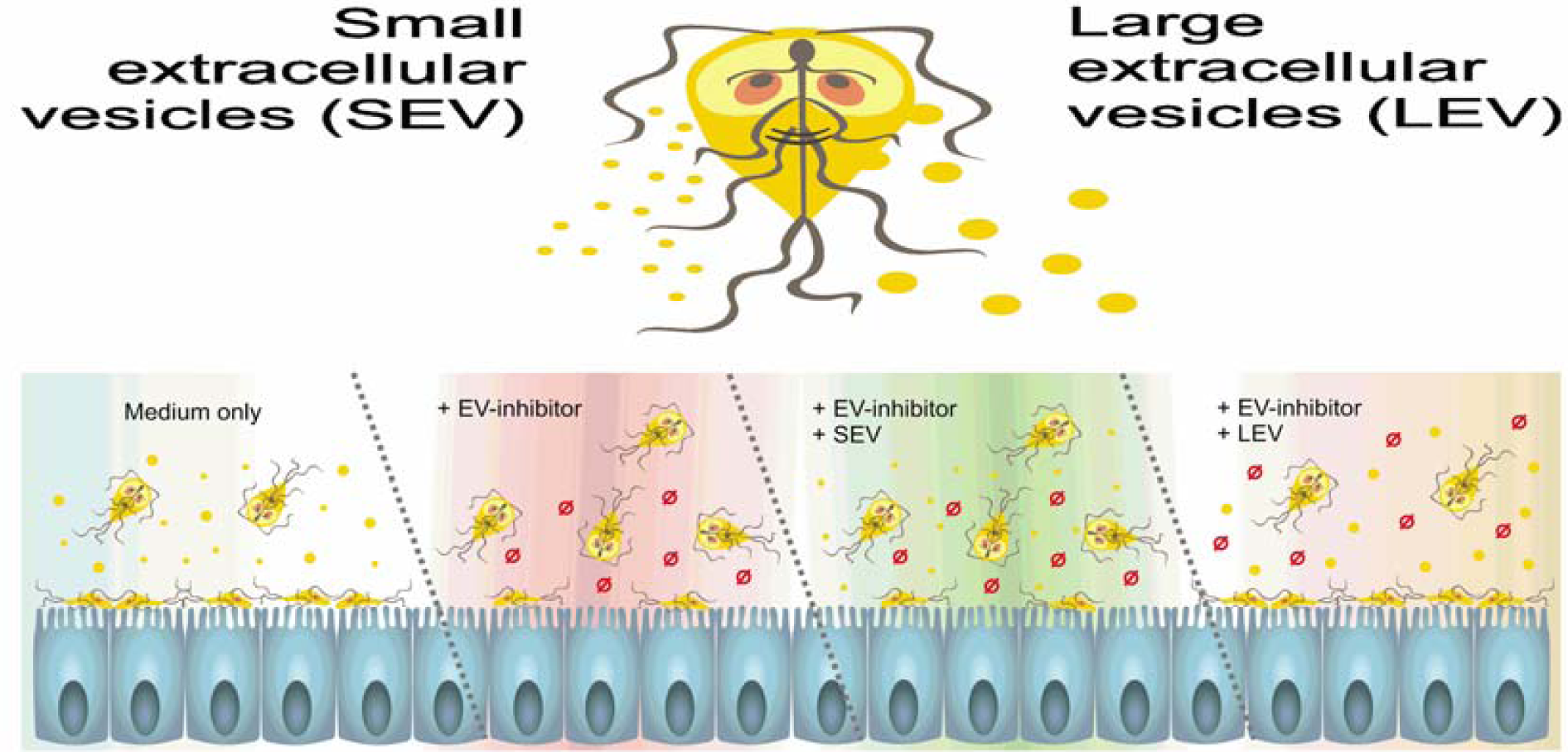
*G. intestinalis* EV populations differentially restore adhesion of trophozoite treated with EV-inhibitors.

EVs from *Giardia* have previously been studied in host-pathogen interactions through proteomic analysis of excretory-secretory products (ESP), including EVs of axenic cultures and cultures of trophozoite interacting with mammalian cells, where proteins related to metabolism were found, without signal peptides on EVs (Ma’ayeh et al 2017). In addition, ESP containing cleavage activity trough cysteine proteases have been identified (de la Mora-de la Mora et al., 2019). The secretome of extracellular protozoa are important for host manipulation and reports indicate a participation of EVs also in other models including *Acanthamoeba castellanii* (Gonçalvez et al., 2018), *Trypanosoma brucei rhodeniense* (Geiger et al., 2010) and *Trichomonas vaginalis* (Twu et al., 2013).

While the majority of *Giardia* infected individuals are asymptomatic, giardiasis is a major contributor to malnutrition and growth impairment in children from developing countries (Fink & Singer, 2017). Additionally, the disease may also last for a long term as a chronic infection. For still unknown reasons, chronic development is associated with sequelae such as malnutrition, inflammatory manifestations, irritable bowel syndrome and extra-intestinal outcomes, including arthritis and food allergy (Hanevik et al, 2014; Bartelt & Sartor, 2015). Adaptive immunity is fundamental to parasite clearance, but leukocytes related with memory such as CD8^+^ T also contribute to chronic inflammation (Scott et al, 2004). Since trophozoites secrete many virulence factors, which contribute to recurrent cycles of reinfection, their resistance to the most used drugs (metronidazole, albendazole) has already been described (Upcroft, 1998; Arguelo-Garcia et al., 2015, Ansell et al., 2015) and the only licensed vaccine is solely used for veterinary practice. Therefore, it is important to identify and study novel clinical strategies that can lead to host recovery. Since PAD-inhibitor Cl-amidine and CBD can effectively decrease parasitic EV release, which contributes to parasite persistence into the small intestine, they may pose as novel therapeutic candidate agents for cases of chronic giardiasis. While important roles for arginine deiminase have previously been established in *Giardia* (Stadelmann et al., 2013; Trejo-Soto et al, 2016; Munoz-Cruz, 2018), a link to EV release has not been made before in *Giardia.* Cannabinoids have previously been linked to inhibition of parasite invasion and immunosuppression of trypanosomiasis (Nok et al., 1994; Croxford et al., 2005) as well as acting as anti-helmitics (Roulette et al., 2016), but their effects on *Giardia* have hitherto not been investigated. Hovewer, GiADI has PAD activity (Touz et al., 2008; Vranych et al., 2014). For example citrullination of the antigenic Variable Surface Proteins (VSPs) through arginine residues of its cytoplasmic tail results in lower antigenic switch, interfering in trophozoite fitness due to cytotoxic antibody activity (Touz et al., 2008). Furthermore, the incomplete conception of *G. intestinalis* pathophysiology and co-pathogen interactions needs to be improved to clarify when the protozoa begins the chronic outcome.

The field of EV research is still rapidly growing, with characterization of functions of subpopulations gaining increased attention. The complex function of LEVs revealed here in *Giardia*, suggests that their influence on phenotype could be even more diverse than those of SEVs (Tckach et al, 2018). No biomarkers were considered in the present study, since both EV populations are enriched mixtures of vesicles that fail to contain any unique marker (Kalra et al., 2013; Vader et al., 2016) and protozoa cells may have different sets of markers in their genome (Gonçalves et al., 2018; Ramirez et al., 2018).

Properties related to different functions of LEV have been studied in non-infectious models. For example, LEVs derived from cancer prostate cells contain substantially more large size ds DNA than SEVs (Vagner et al., 2018). LEVs (microvesicles) derived from platelets were also associated with polymorphonuclear leucocytes increase in adhesion (Fujimi et al., 2002). On the other hand, properties related to cellular adhesion for SEVs isolated from two cancer cell lines have also been identified while the same was not detected for LEVs (Jimenez et al., 2019).

Two subsets of EVs have previously been identified in a parasite model (Cwiklinski et al., 2015), where LEVs contained cargo related to digestion (cathepsin L1 zymogen), while proteomic and functional analyses identified membrane structure components and immunomodulation factors in SEVs. The pan-PAD-inhibitor Cl-amidine has previously been described as a potent EV- inhibitor, compared to a range of other compounds, in various cancer cells (Kosgodage et al., 2017, Kosgodage et al., 2018a), as well as to affect EV-mediated microRNA export (Kosgodage et al., 2019). Previous work has also suggested that PAD-inhibitors can be strategically used to sensitize cancer cells to chemotherapy (Kholia et al., 2015; Kosgodage et al., 2017). The EV-modulatory functions of CBD were recently revealed, and it has been found to be a more potent EV inhibitor than Cl-amidine in some cancers, also to have chemosensiting effects and shows selective inhibition on smaller or larger EVs according to cancer type (Kosgodage et al., 2018b).

In the current study, both PAD-inhibitor and CBD were able to decrease adhesion of *Giardia* to mammalian cells, similar to as our group previously observed with the cholesterol-chelating agent methyl-β-cyclodextrin treatment (Evans-Osses et al., 2017). For our protozoa model in the current study, Cl-amidine regulated LEVs specifically.

The mechanisms of EV biogenesis are still unclear and under ongoing investigation. Inhibition of specific proteins (TSG 101, STAM1 and HRS) has been shown to induce a decrease in exosome secretion, as well as protein content (Colombo et al., 2013). Silencing of specific effectors of the Rab family GTPases on HeLa B6H4 also inhibits exosome secretion, but without modifying cargo or morphology (Ostrowski et al., 2010). Roles for scramblase have been investigated in EV production in *Cryptococcus gatti* including effects on EV size and RNA cargo changes (Reis et al., 2019).

Biomolecular sorting in *G. intestinalis* EVs still needs to be fully understood, including that of acid nucleic populations. As the protozoa lacks a typical endosomal pathway, it is not known if cargo is targeted by post-translational modifications for vesicular secretion. Following the same perspective, not much is understood about RNAi pathways for *Giardia*, although is assumed that a post-translational machinery is necessary to synchronize both transcriptionally active nuclei of the parasite (Adam, 2001; Prucca et al., 2008). A post-transcriptional system related to antigenic switching has been described (Prucca et al., 2016). Serradell et al (2016), produced trophozoite clones expressing a collection of variable-specific surface proteins in plasma membrane through the knockdown of Dicer. Known miRNAs in *Giardia* are derived from hairpins codified in snoRNA (Saraiya et al., 2008; Saraiya et al., 2014). EV could be taken up and influence the host cell. Studies involving uptake from EV serum-derived were already reported for bacterial models (van Bergenhenegouwen et al., 2014; Hiemstra et al., 2014; Yu et al.,2019). However, to add a layer of complexity to the issue, uptake involving eukaryotic cells does not seems unspecific. For example, exosomes derived from oligodendrocytes were internalized (micropinocytosis) by microglia cells, but not by astrocytes (Fitzner et al., 2011). Ofir-Birin et al (2018) reported that *Plasmodium falciparum* derived EVs were processed in host monocytes, as well as their DNA cargo. Szempruch et al (2016) demonstrated that *Trypanosoma brucei* EVs are taken up by mammalian erythrocytes, resulting in anemia. This indicates that EVs accumulate on specific tissues, and that EV uptake can manipulate the host response.

It is possible that small RNAs involved in regulatory functions contained on *G. intestinalis* EVs could silence genes of intestinal immune response, permitting trophozoite adhesion and the low-course inflammation, characteristic of giardiasis. Lambertz et al (2015) observed delivery of small RNAs derived from *Leishmania* exosomes into macrophages, possibly regulating host-pathogen interactions.

## Conclusion

Our results suggest that the two EV populations identified in *G. intestinalis* so far, LEVs and SEVs, have distinct functions in the phenotype of this pathogen and can be selectively modulated using PAD-inhibitor and CBD. Luminal ecology needs to be investigated to completely understand the dual role of *Giardia* as a pathogen/commensal. Since adhesion in the epithelial intestine is fundamental to protozoa fitness, and LEVs clearly aid this process, the use of selective EV–inhibitors, such as Cl-amidine, can be used to interfere with EV secretion, allowing novel strategies to enhance the control of giardiasis.

## Acknowledgements

We would like to thank Dr. Wanderson Da Rocha for sharing his laboratory at the Universidade Federal do Parana. Finally, this study has received support from FIOCRUZ, CNPq, CAPES and Programa Basico de Parasitologia AUXPE 2041/2011 (CAPES), Brazil. M.R is currently fellow from CNPq-Brazil.

